# A model of schema learning based on biological dimensionality reduction during sleep

**DOI:** 10.64898/2026.05.27.728344

**Authors:** Kensuke Yoshida, Genki Shimizu, Yuri Kinoshita, Kaoru Inokuchi, Taro Toyoizumi

## Abstract

Reorganizing learned knowledge into generalized representations and transferring it to future learning are essential aspects of cognition, often described as schema formation and use. However, the computational and circuit mechanisms underlying these processes remain unclear. Here, we propose a theoretical model in which schemas emerge through the formation and alignment of low-dimensional neural representations. In this model, high-dimensional input patterns are reorganized into low-dimensional manifolds through replay-driven Hebbian nonlinear dimensionality reduction. Manifold alignment simultaneously maps representations with shared task structure onto a common format, enabling downstream readout circuits to be reused across tasks. The model captures three core features of schema learning: rapid learning of similar tasks by reusing low-dimensional representations from prior experience, sleep-dependent generalization to unobserved relationships in transitive inference, and compositional recombination of schemas to solve novel tasks. Together, these results suggest a potential neural mechanism for forming, aligning, and recombining low-dimensional schemas to support future learning.

## Introduction

The brain extracts abstract structures from limited experience and uses them for future learning. Such reusable knowledge structures are often referred to as schemas [1–4]. Schema formation and subsequent use may proceed not only during experience itself but also after experience, particularly during sleep and quiet rest [5–11], when replay of awake neural activity patterns [12, 13] has been proposed as a candidate mechanism. However, the mechanisms by which experience is transformed into reusable abstract representations and subsequently used across different situations remain incompletely understood.

Schemas can support future learning through several forms of generalization. First, schemas support transfer across tasks, allowing latent structures acquired in one task to be used in another [14–19]. For example, in reversal learning tasks in which the rewarded choice switches regularly, learning becomes progressively faster when different problems share a common task structure, even when the specific choices differ [14]. Moreover, such task structures shared across distinct problems have been shown to be represented abstractly in the prefrontal cortex [14, 20, 21]. These findings suggest that the brain extracts latent structures common to tasks, rather than merely learning stimulus–reward associations, and reuses them to solve new problems.

Second, schemas support generalization over relationships that were never directly experienced. By abstracting individual experiences into a compact relational structure, a schema may allow unknown relationships to be inferred from that structure [1, 22–24]. Transitive inference provides a canonical example: an unexperienced relation, such as *A > C*, can be inferred from learned relations such as *A > B* and *B > C* [25–27]. This ability requires extracting an ordinal structure beyond pairwise associations and has been reported to be facilitated by sleep [28, 29], suggesting that offline processes may help integrate individual experiences into relational structures that support generalization.

Beyond the use of a single schema for transfer or inference, flexible behavior may require multiple schemas or computational motifs to be combined. Compositional computation refers to the ability to flexibly combine independently learned knowledge or computational elements according to context, thereby adapting to novel tasks or behavioral demands [30, 31]. Replay may support such recombination by reactivating learned elements of experience [30]. Consistent with this idea, a recent study in monkeys showed that, when animals switched among multiple compositionally related tasks, prefrontal activity used a low-dimensional neural subspace shared across tasks to support task performance [32]. These findings suggest that the brain may achieve flexible behavior not by constructing entirely independent representations for each task, but by reusing and recombining common computational elements within a low-dimensional space [32–34].

Together, these findings suggest that seemingly distinct forms of generalization may arise from low-dimensional structural representations extracted from experience [17, 20, 33, 35–38]. In this view, schema formation involves extracting compact relational structure from experience, whereas schema-guided transfer, inference, and composition reflect several ways in which such structure is reused across future learning problems. However, how such representations are formed, used, and recombined to support future learning remains unclear.

Here, we propose a neural circuit model of schema formation, alignment, and composition that combines nonlinear dimensionality reduction and manifold alignment based on biologically plausible synaptic plasticity [39–41]. In this framework, replay transforms high-dimensional representations of experience into low-dimensional schemas that capture relational structure; manifold alignment maps such schemas onto a common format, enabling transfer across tasks and reuse of downstream value-learning circuits; and compositional learning arises from recombining multiple independently acquired schemas. Using this model, we show that transfer between related tasks, the acquisition of transitive inference, and compositional learning that combines transitive structure with reversal learning can be explained as the acquisition, alignment, and composition of common lowdimensional representations. Together, these results suggest a potential computational mechanism by which neural circuits may form reusable low-dimensional schemas from experience and leverage them to support future learning.

## Results

### Model of schema learning

We first constructed a neural circuit model to examine how schemas can be acquired from task experience and subsequently reused for future learning. To formalize the distinction between rote memorization and schema-based generalization, we introduced two conceptually distinct pathways: a direct pathway for rote, task-specific value learning and an indirect pathway for schema-based value learning (Fig. 1a). The direct pathway learned state–value associations independently for each experienced state, whereas the indirect pathway learned values through the schema representation. Here, we implemented the schema representation as a low-dimensional representation that captures the relational structure among experienced states. We used this two-pathway model to examine how this low-dimensional representation enables schema formation, reuse, and transfer across tasks, beyond rote state-specific learning.

**Figure 1:**
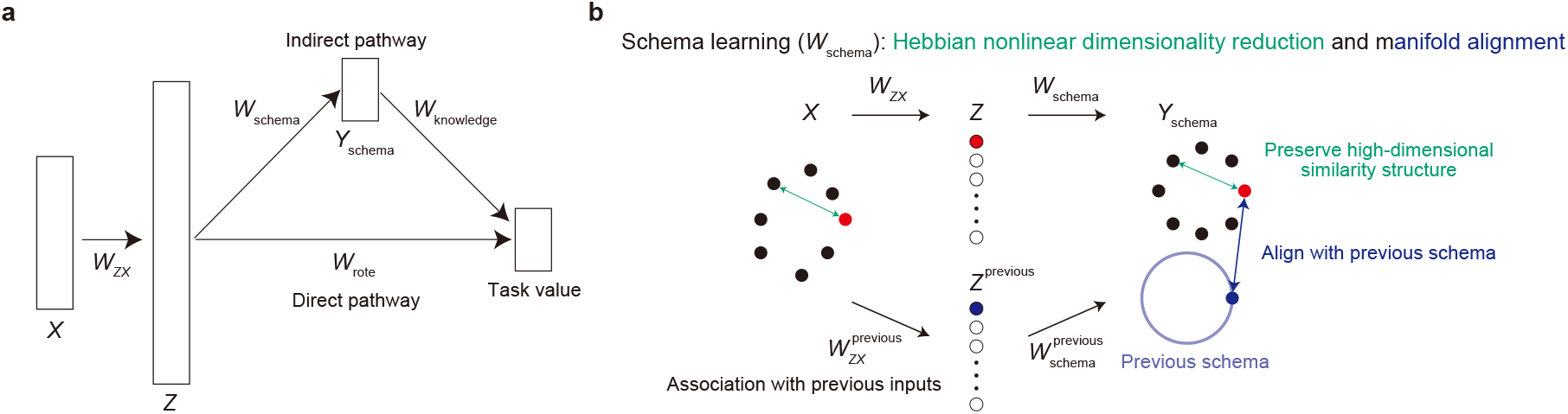
Overview of the model. (a) Schematic of the model. High-dimensional inputs *X* are projected through fixed weights *W*_*ZX*_ and a sharp softmax nonlinearity to yield an intermediate representation *Z*. In the direct pathway, task values are learned from *Z* through the rote readout weights *W*_rote_. In the schema pathway, *Z* is transformed into a low-dimensional schema representation *Y*_schema_ through *W*_schema_, and task values are learned through *W*_knowledge_. The readout weights are updated to reduce reward prediction errors. During sleep, *W*_schema_ is updated by Hebbian nonlinear dimensionality reduction and, when a previous schema is available, by manifold alignment. (b) Illustration of Hebbian nonlinear dimensionality reduction and manifold alignment. Hebbian nonlinear dimensionality reduction preserves neighborhood relationships among high-dimensional inputs in the low-dimensional schema space, whereas manifold alignment pulls current schema coordinates toward reference coordinates computed from a previously learned schema. This allows a new task to acquire a schema that preserves its own local structure while sharing the abstract geometry of a previous task.

Both pathways shared the initial transformation from the high-dimensional state input *X* to an intermediate representation *Z*. This transformation was implemented as a fixed projection, defined by the synaptic weight matrix *W*_*ZX*_, followed by a sharp softmax normalization. The weight matrix *W*_*ZX*_ was constructed from normalized experienced state patterns, so that each intermediate-layer unit corresponded to a stored state pattern.

Thus, *W*_*ZX*_ served as an associative memory: each input activated intermediate-layer units according to its similarity to stored patterns. Because the inverse temperature of the softmax was set high, this transformation approximated a winner-take-all mapping, in which each input predominantly activated the unit corresponding to the most similar stored state.

The direct pathway supported non-schema-based value learning. In this pathway, the task-value readout, with weights *W*_rote_, learned values from *Z* by reducing the prediction error between the reward target and the predicted value. Because *Z* was approximately winner-take-all, value updates were largely local to the activated state representation. This pathway therefore corresponds to rote memorization, in which state–value associations are acquired independently for each input.

The schema pathway provided an additional low-dimensional representation for value learning. The intermediate representation *Z* was transformed into a schema representation *Y*_schema_ through the schema weight matrix *W*_schema_. The task-value readout, with weights *W*_knowledge_, was trained by the same rule as in the direct pathway to reduce the reward prediction error. The critical difference was therefore the structure of the representation available to the readout. Before schema learning, value prediction depended primarily on the state-specific representation *Z*; after schema learning, the readout could additionally exploit the relational structure encoded in *Y*_schema_.

The key component of the model was the learning of *W*_schema_. Unlike the task-value readout weights, *W*_schema_ was not updated by reward prediction errors, but was instead constrained by two requirements. First, the schema should represent the relational structure of the current task by organizing high-dimensional state representations into a lowdimensional manifold. Second, when a related task had already been experienced, the new schema should be associated with the previously acquired schema so that common task structure could be reused. In the present model, these two requirements were implemented as two complementary objectives: Hebbian nonlinear dimensionality reduction [39] for extracting the structure of the current task and manifold alignment for linking the new schema to a previously learned one.

*W*_schema_ was updated to produce a low-dimensional representation that preserved the neighborhood structure of the original high-dimensional state space (Fig. 1b). We implemented this process using Hebbian nonlinear dimensionality reduction, a biologically plausible nonlinear dimensionality reduction rule proposed in previous work [39]. Intuitively, Hebbian nonlinear dimensionality reduction follows the same principle as t-SNE [42]: it organizes low-dimensional representations so that pairwise similarities in the low-dimensional space match pairwise similarities in the original high-dimensional space. Hebbian nonlinear dimensionality reduction provides a biologically plausible implementation of this optimization through repeated presentation of input patterns and a three-factor Hebbian plasticity rule [40, 41]. We assigned this learning process to sleep because sleep provides a natural offline regime for replay-driven reorganization, in which previously experienced state representations can be reactivated repeatedly in the absence of ongoing task demands [5–7, 11]. Thus, sleep-dependent updates of *W*_schema_ allowed the model to extract the relational structure of the current task independently of reward prediction errors.

To model transfer across related tasks, we introduced a manifold alignment mechanism that used a previously learned schema as a reference (Fig. 1b). For each state in a new task, the model computed a reference schema coordinate by projecting the new input through the previous task’s prototype-based intermediate layer and then through the previous schema weights. This procedure provided a soft correspondence between new-task states and positions on the old schema manifold, without requiring explicit state matching. During sleep in the new task, *W*_schema_ was updated by the Hebbian nonlinear dimensionality reduction objective together with an alignment loss that pulled current schema coordinates toward these reference coordinates. Thus, the new schema preserved local relationships among current-task states while inheriting the abstract geometry of the previously learned schema.

Importantly, this alignment mechanism does not require sensory inputs to be identical across tasks. Even when sensory representations are largely orthogonal, tasks may share weak common components, such as action structure, reward contingencies, or other task-relevant variables. Because alignment is performed in a low-dimensional schema space, such weak shared components can provide sufficient constraints: there are fewer degrees of freedom for relative rotations between representations than in the original high-dimensional sensory space. The model can therefore use these shared components to establish a soft correspondence with the previous schema manifold, enabling transfer through partial overlap in behaviorally relevant structure rather than through direct matching of sensory states. In the following sections, we test this idea in tasks in which sensory inputs differ across conditions but the underlying action or reward structure provides a common axis for alignment.

The model constructed here formalizes schema learning as the offline construction of a reusable low-dimensional representation. The direct pathway supports rote value learning when no schema is available, whereas the schema pathway allows value learning to exploit relationships among states once a structured representation has been formed. We therefore asked whether this mechanism could account for faster acquisition within a task and transfer across tasks with different sensory inputs but shared abstract structure.

### Model enables transfer learning between spatial tasks

We first tested the model in a spatial alternation task, similar to a previous experimental study [15], in which the agent had to alternate left and right choices across successive trials (Fig. 2a). We used this task to examine whether sleep-dependent schema formation could extract a low-dimensional representation of the task structure and facilitate subsequent value learning.

**Figure 2:**
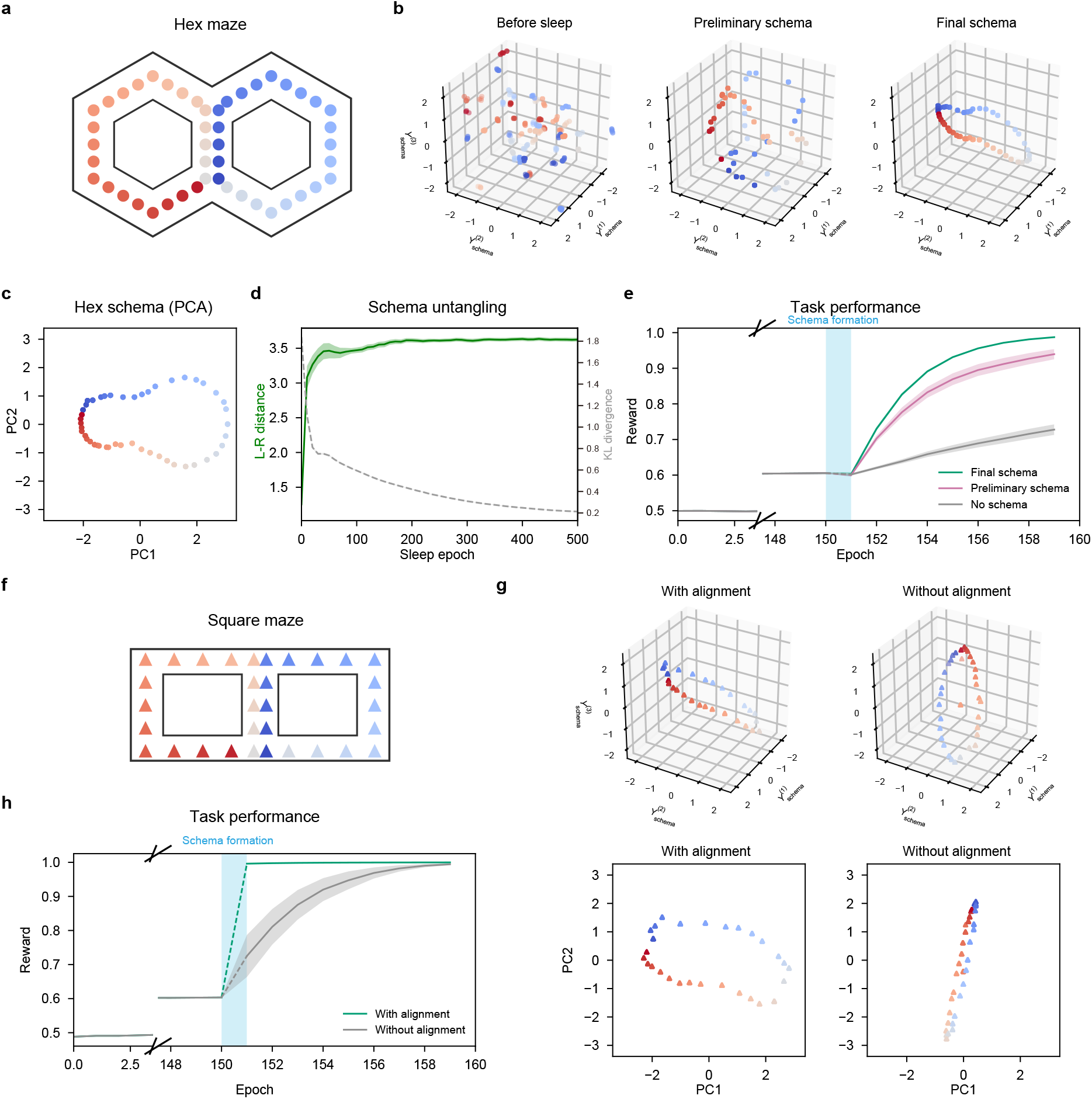
Schema learning accelerates and transfers maze learning. (a) Schematic of the hexmaze spatial alternation task. Reward was obtained by alternating left–right choices across successive trials. (b) Three-dimensional schema representations before sleep, after preliminary schema formation, and after final schema formation. Points are colored according to maze position. (c) Two-dimensional PCA projection of the final hex-maze schema. (d) Left–right schema distance and KL divergence across sleep epochs (mean ± SEM, *n* = 25 simulations). The left–right schema distance quantifies separation between states associated with left versus right trajectories in schema space, and the KL divergence corresponds to the Hebbian nonlinear dimensionality reduction objective. (e) Time course of task performance in the hex maze (mean ± SEM, *n* = 25 simulations). The vertical blue band marks the sleep period during which the schema was formed. (f) Schematic of the square-maze spatial alternation task. As in the hex-maze task, reward was obtained by alternating left–right choices across successive trials. (g) Three-dimensional square-maze schema representations and their projections onto the first two principal components defined from the final hex-maze schema in (c). With alignment, the square-maze schema recapitulated the abstract geometry learned in the hex maze; without alignment, the schema formed a different orientation. (h) Transfer performance across wake epochs in the square maze (mean ± SEM, *n* = 25 simulations). Alignment allowed the previously learned schema-based readout to be reused immediately.

The network received high-dimensional state inputs that encoded the agent’s recent spatial trajectory together with task-relevant auxiliary variables. The spatial component was constructed as a temporally smoothed representation of the agent’s position history, allowing each input to contain information about the recent path taken by the agent. The auxiliary variables encoded the selected action, namely whether the agent turned left or right, and the reward outcome obtained after passing through the choice points (see Methods).

During initial wakefulness, the schema pathway was absent, and the task-value readout learned only from the fixed intermediate representation. Because the intermediate representation did not explicitly encode the relational structure of the task, learning progressed slowly (Fig. 2e). Sleep was then inserted as an offline period in which previously experienced state representations were replayed, and the schema weights were updated by Hebbian nonlinear dimensionality reduction. Across sleep, the initially unstructured schema representation gradually became organized according to the latent left–right alternation structure of the task (Fig. 2b,c). In the final-schema condition, the learned manifold formed a ring-like structure that reflected the periodic structure of the task, in which left- and right-associated states were visited alternately (Fig. 2c). Consistent with this qualitative organization, the objective function for Hebbian nonlinear dimensionality reduction, measured by the KL divergence between high-dimensional and low-dimensional similarity structures, gradually decreased during sleep (Fig. 2d). Moreover, states that should be distinguished for successful alternation, such as states reached after left versus right trajectories, became more clearly separated (Fig. 2b,d). This ring-like representation is consistent with the previously observed low-dimensional CA1 activity patterns in the same task [15]. After sleep, the model using the final schema showed a faster increase in expected reward than models using either a preliminary schema or no schema (Fig. 2e), indicating that the schema formed during sleep enabled rapid value learning by exploiting the relational structure of the task.

We next asked whether the learned schema could transfer to a new spatial task. After learning the original hexagonal maze (Fig. 2a), the model learned a second maze with a different sensory geometry, a square maze (Fig. 2f). The place representations of the two mazes were set to be orthogonal in the input space, so transfer could not be achieved by directly reusing maze-specific sensory features. However, the two tasks shared a limited set of behaviorally relevant auxiliary variables, including left/right turn information and reward outcomes, reflecting their common abstract alternation structure. This setting allowed us to test whether schema transfer can occur when sensory inputs are orthogonal across tasks but a small set of task-relevant variables provides common anchors for alignment.

When schema alignment was applied during sleep in the new maze, the resulting representation was successfully aligned with the left–right organization learned in the original maze (Fig. 2b,c,g). By contrast, without alignment, the schema learned in the new maze did not match the previously acquired structure (Fig. 2b,c,g). Even before the value-readout weights were further adjusted in the new maze, the aligned model showed improved task-value predictions simply by aligning the new schema to the previously learned manifold (Fig. 2h). Thus, alignment alone allowed the model to immediately benefit from the previously acquired schema, before additional task-specific value learning. These results are consistent with previous experimental findings that mice learned a similar maze more rapidly after prior experience and that low-dimensional CA1 activity patterns were aligned between the first and second mazes [15]. These results show that the model can transfer an abstract spatial schema across tasks even when the sensory representations of the environments are different.

Together, the spatial task simulations demonstrate two key functions of the model. First, sleep-dependent Hebbian nonlinear dimensionality reduction extracts a low-dimensional schema that improves value learning within a task. Second, manifold alignment allows this schema to be reused across tasks that share a common relational structure. Thus, schema learning in the model does not merely compress sensory inputs, but constructs an abstract representation that can support transfer learning across different tasks.

### Model enables transfer learning between non-spatial tasks

We next asked whether the same mechanism could support transfer in a non-spatial task, in which the relevant structure was defined not by spatial trajectories but by the temporal organization of choices and rewards. For this purpose, we used a reversal learning task, similar to a previous experimental study [14], in which the currently rewarded choice reversed every 10 trials according to a latent rule (Fig. 3a). Successful behavior in this task requires the model to infer the current rule from recent choice and reward history, rather than simply associating each sensory input with a fixed outcome.

**Figure 3:**
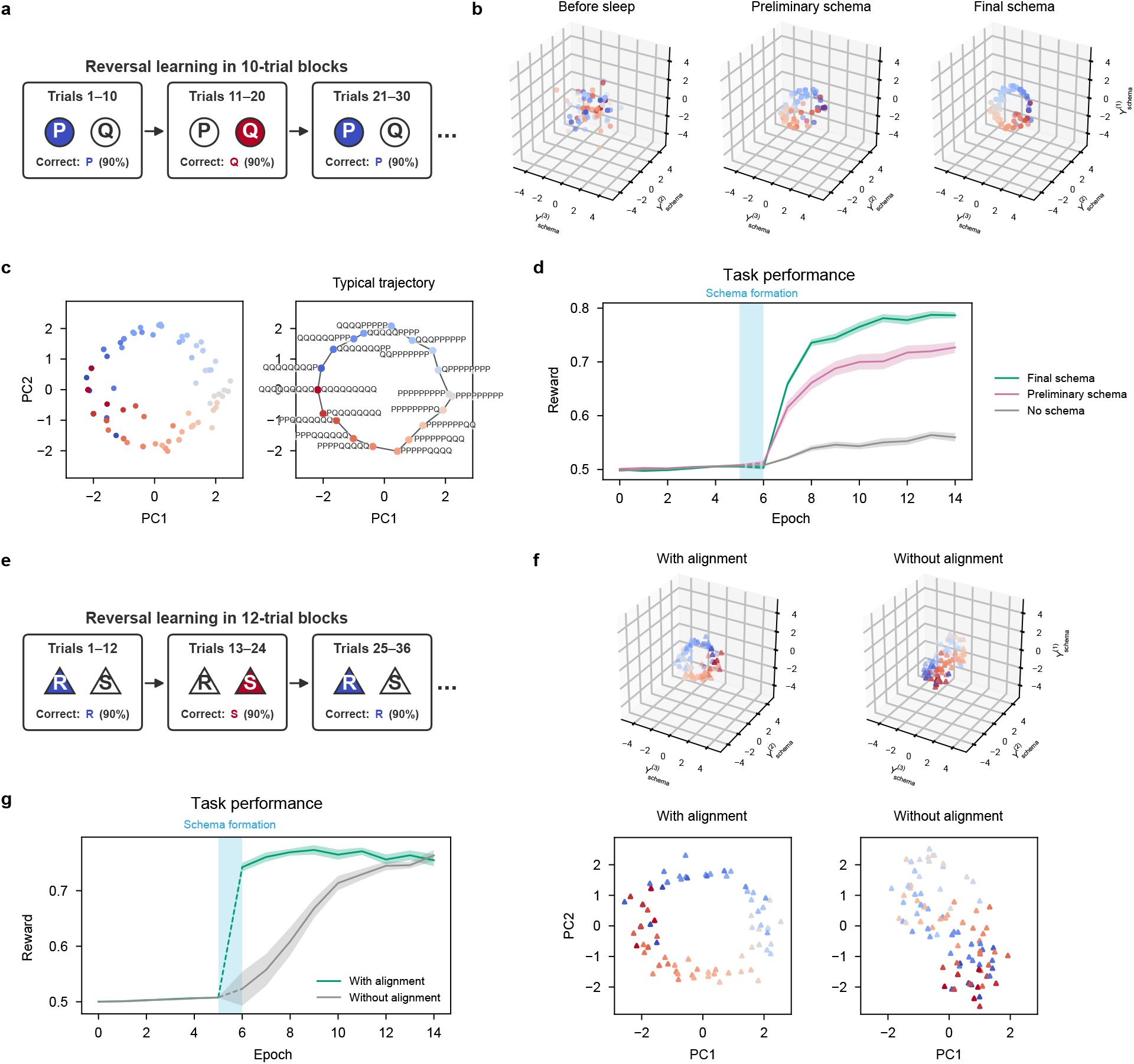
Schema learning accelerates and transfers reversal learning. (a) Schematic of the reversal learning task. The rewarded choice reverses every 10 trials. (b) Three-dimensional schema representations before sleep, after preliminary schema formation, and after final schema formation. Points are colored according to task state. (c) Left, two-dimensional PCA projection of the Task 1 schema in the Final schema condition. Right, projection of a typical sequence of states spanning one rule cycle, illustrating that successive states trace a ring-like trajectory in schema space. (d) Task performance across wake epochs for three conditions (Final schema, Preliminary schema, and No schema; mean ± SEM, *n* = 25 simulations). The vertical blue band marks the sleep period during which the schema was formed. (e) Schematic of a second reversal-learning task with a distinct physical implementation and a different reversal schedule. (f) Three-dimensional schema representations and the corresponding PC1/PC2 projections, with and without alignment. (g) Task 2 performance for the With alignment and Without alignment conditions (mean ± SEM, *n* = 25 simulations). Alignment enabled faster learning by allowing the Task 1 schema-based readout to be reused in Task 2.

The model received high-dimensional inputs that encoded recent correct-choice history together with reward-related auxiliary variables (see Methods). As in the spatial task, the schema pathway was absent before sleep, and value learning depended on the fixed intermediate representation. Sleep was then inserted, during which the schema weights were updated by Hebbian nonlinear dimensionality reduction. Across sleep, the history-dependent states became organized into a low-dimensional schema. The resulting representation captured the cyclic structure of reversal learning: states associated with different phases of the latent rule were arranged along a ring-like manifold in schema space (Fig. 3b,c). After sleep, the model using the final schema learned the reversal task more rapidly than models using either a preliminary schema or no schema (Fig. 3d). This result indicates that the hidden cyclic structure of reversal learning was acquired as a schema and used to guide subsequent task-value learning to exploit the relational structure among history-dependent states.

We then examined whether this non-spatial schema could be reused in a related reversal task. After learning the first reversal task, the model learned a second task with a different reversal period, in which the rewarded choice reversed every 12 trials (Fig. 3e). The sensory inputs of the two tasks were orthogonal, whereas the reward-related auxiliary variables were shared across tasks (see Methods). Thus, this setting allowed us to test whether schema transfer can occur through weak shared cues related to reward contingencies.

When schema alignment was applied during sleep in the second task, the new lowdimensional representation aligned with the schema learned in the first task (Fig. 3b,c,f). Specifically, states in the 12-trial reversal task were mapped onto corresponding phases of the previously learned 10-trial reversal schema. This representational alignment facilitated transfer: even before additional task-value readout learning in the second task, schema alignment alone was sufficient to produce near-complete learning performance (Fig. 3g). These results show that schema transfer can occur in a non-spatial task even when sensory inputs are orthogonal and reversal schedules differ. This parallels the prior behavioral and neural findings [14]: mice learned new reversal-learning problems more rapidly across problems with distinct physical implementations, and prefrontal populations represented task variables in a format that generalized across problems. In the present model, such faster learning arises because manifold alignment maps task-specific inputs onto a shared low-dimensional schema, allowing the same value readout to be reused across problems.

Together, these results suggest that sleep-dependent schema formation and alignment can support transfer over abstract task variables when different experiences share a common relational organization.

### Schema learning facilitates transitive inference

We next tested whether the schema pathway could support not only transfer across tasks with shared structure, but also generalization over relations that were not directly trained. A canonical example of such relational generalization is transitive inference, in which an ordered structure must be inferred from overlapping pairwise experiences. We simulated a task in which items formed a latent hierarchy, but the model was trained only on adjacent pairs (Fig. 4a). Because non-adjacent pairs were never reinforced directly, successful choices on these probe trials required integration across learned relations rather than memorization of individual pair outcomes.

**Figure 4:**
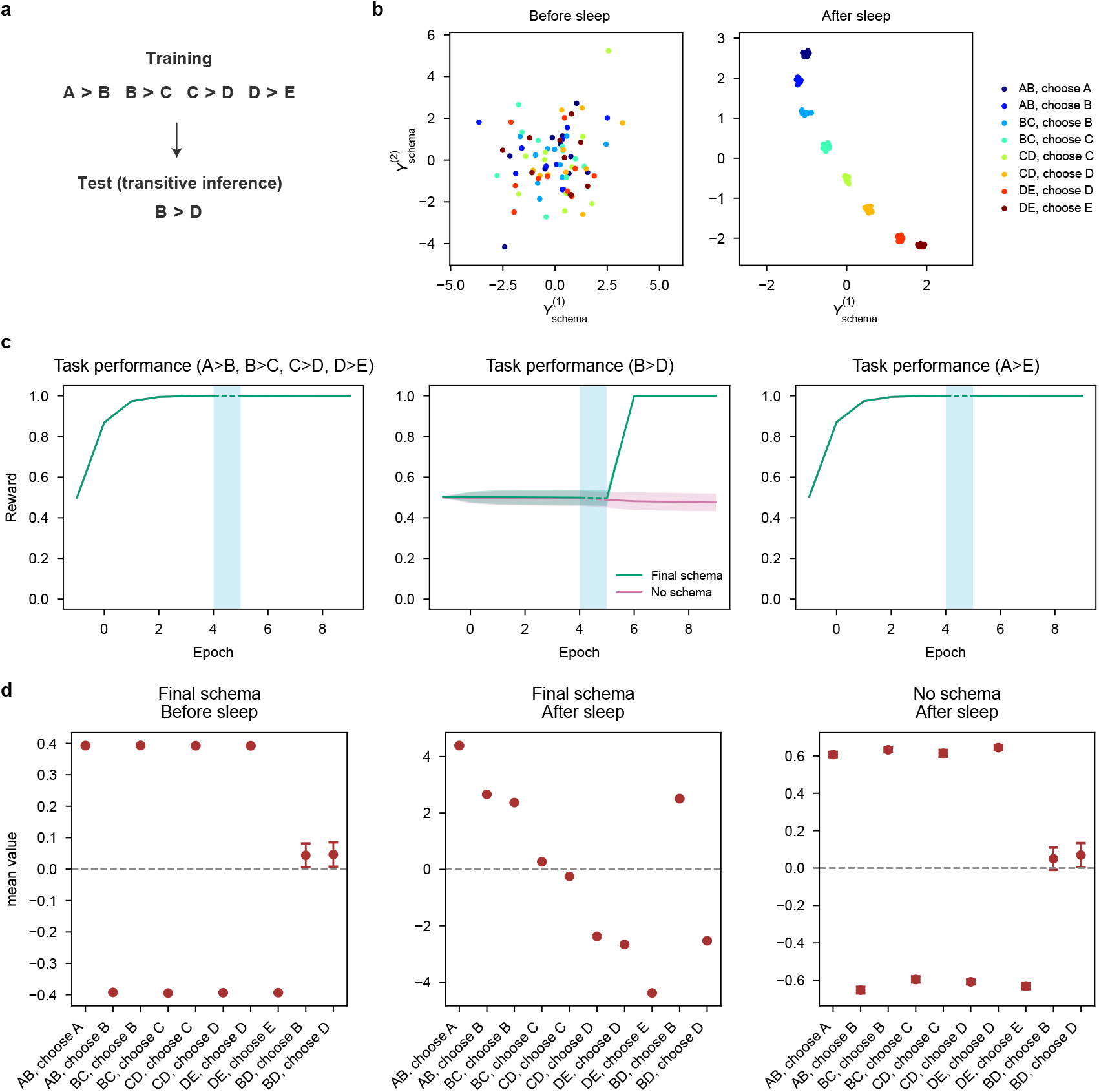
Sleep-dependent schema formation enables transitive inference. (a) Schematic of the transitive inference (TI) task. Items A–E are ordered *A > B > C > D > E*, but only adjacent pairs (AB, BC, CD, DE) were trained. (b) Representative two-dimensional schema representations before and after sleep. After sleep, the schema organized states according to the latent rank order. (c) Task performance for trained adjacent pairs, the non-adjacent BD test pair, and the end-anchor AE pair (mean ± SEM, *n* = 100 simulations). The Final schema condition solved the non-adjacent BD test pair after sleep, whereas the No schema condition remained at chance. (d) Value estimates for trained and non-adjacent pairs (mean ± SEM, *n* = 100 simulations). A value hierarchy supporting transitive inference emerged only in the Final schema condition after sleep.

During wakefulness before schema learning, the task-value readout rapidly acquired the directly trained adjacent comparisons by using the direct pathway (Fig. 4c). However, without a structured schema, the model had little basis for assigning consistent relative values to items that had never been directly compared (Fig. 4d). Sleep-dependent Hebbian nonlinear dimensionality reduction changed the geometry of the item representations. The initially unstructured two-dimensional schema became organized along an ordered axis consistent with the latent hierarchy (Fig. 4b). This sleep-generated relational schema supported transitive inference. The model with the final schema maintained strong performance on adjacent pairs and, critically, showed improved performance for the nonadjacent probe pair *B > D* (Fig. 4c). The task-value representation also became more consistent with the abstract item order: items that were never directly contrasted nevertheless acquired relative values appropriate for their positions in the hierarchy, including the preference *B > D* (Fig. 4d). By contrast, the no-schema condition learned the trained pairwise discriminations but failed to construct a coherent ordering over untrained pairs (Fig. 4d).

This pattern is consistent with experimental findings that sleep selectively improves transitive inference on the non-adjacent pair *B > D* [28]. Additionally, the end-anchor pair *A > E* was already solved before sleep in both the experiment and the model (Fig. 4c), suggesting that this comparison can be solved without constructing the full relational schema. Thus, the model reproduced the sleep-dependent improvement in transitive inference.

These results show that the model can use sleep-dependent schema learning to go beyond directly experienced associations. The schema pathway integrated local pairwise relations into a global low-dimensional structure, allowing the task-value readout to generalize over relationships that were not explicitly reinforced. This provides a mechanistic account of how offline replay could support transitive inference by reorganizing overlapping experiences into an ordered relational representation.

### Model composes independently learned schemas to solve a new task

Finally, we asked whether schemas acquired in different tasks could be composed to solve a new task that required multiple abstract computations. To test this possibility, we constructed a task that combined the transitive-inference hierarchy over items *A*–*E* with a reversal-learning schema that had been acquired independently in a two-item task using items *P* and *Q* (Fig. 5a). Here, *P* and *Q* were distinct from the TI items *A*–*E* and were used only to learn the reversal-learning schema. In the new task, the model was trained on adjacent pairs sampled according to the current hierarchy and tested on non-adjacent pairs. Solving this task required two computations: inferring the ordered relational structure among items and flexibly updating which relational rule was currently rewarded.

**Figure 5:**
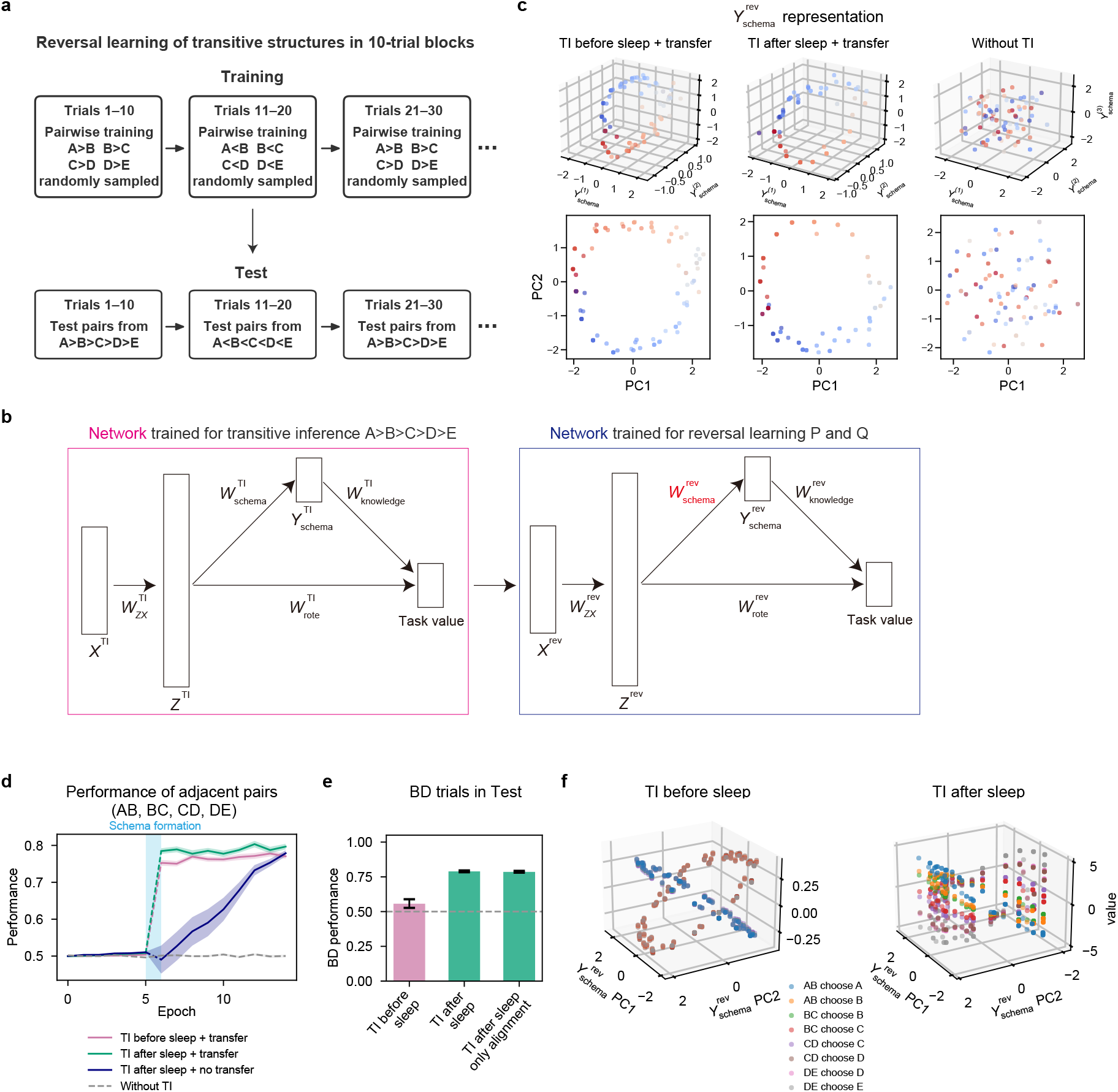
Compositional learning of transitive inference and reversal learning. (a) Schematic of the reversal learning of transitive structures. In each 10-trial block, the rewarded ordering alternates between *A > B > C > D > E* and *A < B < C < D < E*. During training, adjacent pairs are randomly sampled from the current ordering; during test, the model is evaluated on pairs sampled from the same ordering, including non-adjacent pairs. (b) Schematic of the composite network. The transitive-inference network was pretrained on the *A*–*E* hierarchy, whereas the reversal-learning network was pretrained independently in a separate two-item task using *P* and *Q*. The output of the TI network was provided as input to the reversal-learning network. (c) Schema representations for three input conditions of the reversal-learning network: TI-derived values before sleep (“TI before sleep”), TI-derived values after sleep (“TI after sleep”), and raw high-dimensional trial-state history (“Without TI”). Top, three-dimensional schemas obtained at the end of sleep; bottom, the corresponding PC1/PC2 projections. (d) Performance for training adjacent patterns for four conditions: “TI before sleep + transfer”, “TI after sleep + transfer”, “TI after sleep + no transfer”, and “Without TI” (raw high-dimensional trial-state history) (mean ± SEM, *n* = 25 simulations). (e) Performance on the non-adjacent *B > D* trials for the TI-before-sleep, TI-after-sleep, and TI-after-sleep only-alignment conditions (mean ± SEM, *n* = 25). In the only-alignment condition, performance was evaluated immediately after schema alignment, before any additional task-value learning. (f) Compositional value representation. Three-dimensional plots show the reversal-learning state projected onto schema-space PC1 and PC2, with the value computed by composing transitive-inference and reversal-learning representations plotted on the vertical axis, for each of the eight adjacent-pair choices. Results are shown separately for the TI-before-sleep and TI-after-sleep conditions.

We asked whether a model that had separately learned transitive inference and reversal learning could combine these components to solve this new task more efficiently (Fig. 5b). In one network, the model had learned the transitive hierarchy *A > B > C > D > E*. In another network, the model had learned history-dependent reversal learning between two items, *P* and *Q*. We then constructed a composite network in which the output of the transitive-inference network was provided as input to the reversal-learning network (Fig. 5b). In this composite model, the transitive-inference network was kept fixed, and learning in the new task occurred in the reversal-learning network, including updates of 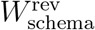,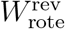, and 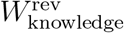. Using this architecture, we examined whether the model could solve a task in which the rewarded relational rule alternated between *A > B > C > D > E* and *A < B < C < D < E* (Fig. 5a).

We compared three conditions that differed in the state of the transitive-inference network. In the first condition, transitive inference had been completed after sleep-dependent schema formation, corresponding to the after-sleep condition in Fig. 4. In the second condition, the adjacent training pairs had been learned, but transitive inference had not yet emerged, corresponding to the before-sleep condition in Fig. 4. In the third condition, no transitive-inference learning had occurred, and the reversal-learning network received only raw high-dimensional trial-state history without a structured TI-derived representation. We further compared two conditions in the reversal-learning network: with or without alignment to the previously learned reversal schema.

When raw trial-state history was provided without the transitive-inference network, the model failed to form a clear reversal-learning structure in 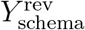 (Fig. 5c). This is likely because the recent history contained intermixed experiences from multiple adjacentpair problems, such as AB, BC, CD, and DE, without an explicit representation of their shared relational meaning. As a result, the reversal-learning network could not easily discover the common latent structure underlying these experiences, and learning progressed only slowly (Fig. 5d).

By contrast, when the input to the reversal-learning network was derived from the transitive-inference network, the model acquired a ring-like structure characteristic of reversal learning (Fig. 5c). This structure was observed in both the TI-before-sleep and TI-after-sleep conditions, indicating that even partially learned TI-derived representations were sufficient to support the discovery of the reversal-learning dynamics for the trained adjacent pairs. In both cases, reversal learning proceeded much more rapidly than in the condition without TI-derived input (Fig. 5d). Moreover, when the TI-derived representation was aligned to the previously learned reversal-learning schema, learning was especially rapid, indicating that the model could benefit from a reversal-learning computation acquired in a separate task.

Importantly, although both the TI-before-sleep and TI-after-sleep conditions supported learning on the trained adjacent-pair problems, they differed in their ability to generalize. We next evaluated the model on an all-pair test version of the task (Fig. 5a), while freezing the weights learned during adjacent-pair training. In this setting, the model had to apply the learned reversal computation to item combinations that had not been directly trained. Only conditions using the TI-after-sleep values showed improved performance on the non-adjacent *B > D* trials, including the condition evaluated immediately after schema alignment and before any additional task-value learning (Fig. 5e). Thus, sleep-dependent completion of the transitive-inference schema was necessary for generalization to non-adjacent relational choices.

This generalization can be understood from the geometry of the learned representations. After sleep, the composite network represented the values of items A–E as a coherent hierarchy within each reversal-learning state (Fig. 5f). Across reversal states, this hierarchy was systematically transformed, such that the behavioral meaning of the rank order changed with recent history. In other words, the model acquired not only a value hierarchy over items, but also a history-dependent transformation of that hierarchy. This indicates that the network compositionally combined the transitive-inference schema with the reversal-learning schema.

This result illustrates the compositional use of schemas. The transitive-inference schema provided an abstract rank representation, whereas the reversal-learning circuit learned how the behavioral relevance of that representation changed with recent history. Importantly, once the two networks had been trained separately, the model could solve the new compositional task through schema alignment alone, without further adjustment of the task-value readout weights (Fig. 5e). Combining these two components allowed the model to solve a task that required both relational generalization and flexible rule switching. Thus, the model provides a circuit-level mechanism by which the brain could solve novel tasks by aligning and recombining schemas acquired in different experiences. More generally, sleep-generated schemas may serve not only as static representations for value learning, but also as reusable computational variables that can be flexibly recombined with other task demands.

## Discussion

In this study, we proposed a biologically plausible circuit model in which sleep-dependent replay constructs low-dimensional schemas and aligns them with existing schemas so that they can be reused for subsequent value learning. The model has two key components. First, replayed high-dimensional experiences are transformed into low-dimensional schema representations through Hebbian nonlinear dimensionality reduction, allowing the relational structure of a task to be extracted without direct reward-based supervision. Second, when a related task has already been experienced, the newly formed schema is aligned with an existing schema, enabling downstream readout circuits to be reused. Through these mechanisms, the model explained, within a single framework, transfer between spatial tasks [15], transfer between non-spatial reversal-learning tasks [14], sleep-dependent improvement in transitive inference [28,29], and compositional behavior in a task combining transitive inference with reversal learning. Thus, the model provides a unified account of how offline replay transforms experiences into abstract formats that can support flexible behavior [11].

The model also provides a computational interpretation of generative replay and hypothesis testing. Previous work has suggested that replay in hippocampal–prefrontal circuits may support compositional inference by generating candidate possibilities and evaluating them against task demands [30, 31]. In the present model, the schema representation *Y*_schema_ can be interpreted as such a candidate hypothesis. During sleep, replay generates a possible low-dimensional organization of the current experience. The subsequent alignment process then tests whether this candidate organization can be mapped onto an existing schema and whether a previously learned readout can be reused. From this perspective, schema learning is not merely a compression of experience, but a process of hypothesis generation and evaluation. In other words, the brain may search for a representational coordinate system in which a new task can be interpreted using computations that are already available.

This interpretation gives a functional role to the coexistence of recent- and prior-memory replay. Recent work has shown that NREM sleep contains microstructurally distinct substates in which recent and prior memories are preferentially replayed [43]. The present model suggests that such temporal organization may be useful for schema alignment. Recent-memory replay provides samples from the newly experienced task, whereas prior-memory replay provides reference samples from previously learned schemas. If these two replay streams occur alternately or in a coordinated manner, the circuit can compare the current manifold with past manifolds and adjust the new representation so that it preserves relationships within the current task while aligning with computational axes that were useful in the past. Thus, the model predicts that disrupting the temporal coordination between recent- and prior-memory replay should impair transfer and schema integration more strongly than simple memory retention.

A central advantage of low-dimensional schema representations is that they constrain the space of possible transformations [35, 44, 45]. In high-dimensional representations, task-relevant variables can in principle rotate, deform, or recombine along many degrees of freedom, making it difficult to identify mappings that allow previously learned readout circuits to be reused. By contrast, when experience is organized into a compact low-dimensional manifold, the relevant transformations are restricted to a smaller set of operations on the latent structure. In our model, this restriction allowed low-dimensional schemas to be aligned using only a small number of anchor points (Figs. 2, 3, and 5). Building on this alignment, the composite network in Fig. 5 showed that a transitive-inference circuit could be linked to a reversal-learning circuit, allowing relational ordering and reversal learning to be combined in a novel task. Thus, dimensionality reduction may support not only the representation of latent structure [46, 47], but also the alignment and composition of multiple schemas for flexible reuse.

This idea naturally connects with recent evidence that the brain can construct compositional tasks using shared neural subspaces [32]. In that study, task-relevant variables were represented in neural subspaces that were reused across multiple tasks. The present model provides one possible learning mechanism for how such shared subspaces could be constructed and linked across experiences. The transitive-inference schema supplied an abstract rank variable, whereas the reversal-learning readout supplied a history-dependent rule that changed the behavioral meaning of that variable. By combining these components, the model solved a new task that required both relational inference and flexible rule switching. Thus, schemas in the present model are not static summaries of memory, but reusable computational variables that can be recombined with other circuits.

The model makes several experimentally testable predictions. First, sleep-dependent improvements in schema-based tasks should be accompanied by a gradual reduction in the dimensionality of task representations and by increased alignment between new and old task manifolds. Second, the degree of alignment between recent- and prior-memory replay contents should predict the degree of transfer across tasks. Third, selective disruption of periods dominated by recent-memory replay should impair the formation of new schemas, whereas disruption of periods dominated by prior-memory replay should impair alignment with existing schemas.

Several questions remain open. First, how does the brain search for a schema that is appropriate for a new task? In the present model, candidate schemas are effectively tested one after another, but biological systems may use a more efficient search strategy. For example, partial similarity between the current experience and existing schemas may bias which schemas are retrieved and tested, or replay itself may prioritize past schemas that share local relational structure with the new task. Second, the distinct roles of NREM and REM sleep remain to be clarified. In transitive inference [25,29], for example, both NREM and REM sleep have been shown to be necessary [28]. One possibility is that NREM sleep preferentially supports the extraction and stabilization of relational structure through replay, whereas REM sleep may support broader recombination, integration, or evaluation of candidate schemas. Incorporating these sleep-stage-specific mechanisms into the model will be an important direction for future work. Third, it remains unclear what determines the transition from rote memorization of individual experiences to schema-based understanding. A previous study showed that transitive inference emerged only after a randomized training phase, in which two learned premise pairs were trained each day but the combination of pairs varied across days [28]. This finding suggests that schema formation may require conditions that force separately learned relationships to be integrated into a common relational structure, rather than simply strengthening each pairwise association. Future work should clarify what signals trigger this transition in biological circuits.

In summary, this study proposed a model in which sleep-dependent replay constructs low-dimensional schemas and aligns them with existing schemas, thereby enabling the reuse of downstream readout circuits. This framework explains transfer learning, transitive inference, and compositional computation through a single circuit principle. More broadly, the model suggests that one major function of replay is to search over lowdimensional hypotheses about the structure of experience, evaluate their compatibility with existing schemas, and transform memories into reusable computational building blocks.

## Methods

### Model architecture

We used a neural circuit model consisting of an input layer *X*, an intermediate layer *Z*, a schema representation layer *Y*_schema_, and a task-value readout (Fig. 1a). Throughout, lowercase boldface symbols, such as ***x***_*i*_ and ***z***_*i*_, denote vectors for sample *i*, corresponding to the layer-level variables, such as *X* and *Z*, introduced in the main text. For an input state ***x***_*i*_, activity in the intermediate layer was given by

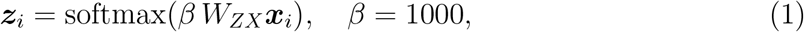

where the synaptic matrix *W*_*ZX*_ was fixed. The rows of *W*_*ZX*_ were normalized prototype patterns constructed from representative task experience. The large inverse temperature made ***z***_*i*_ nearly winner-take-all, so that the intermediate layer acted as a pattern-separated representation of experienced states.

The schema pathway projected ***z***_*i*_ into a low-dimensional schema representation through a plastic synaptic matrix:

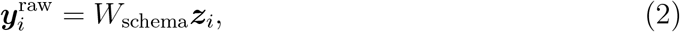

where *W*_schema_ was a plastic synaptic weight matrix. Before being used by the task-value readout, the schema activity was centered and normalized:

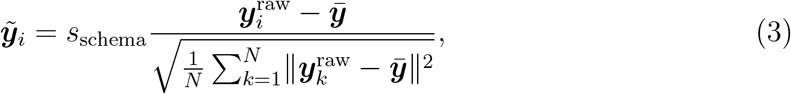

where 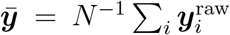 and *s*_schema_ = 2. The normalized vector 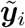 was used as the activity of the schema representation layer *Y*_schema_ for sample *i*.

The task-value readout contained two components. The direct pathway, with weights *W*_rote_, read out value from the intermediate representation ***z***_*i*_ and supported task-specific memorization. The schema-dependent pathway, with weights *W*_knowledge_, read out value from the schema representation 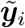 and allowed value learning to exploit relational structure extracted during sleep. The predicted value was

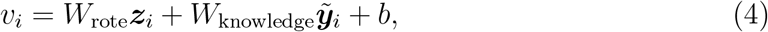

where *b* is a bias term.

### Task-value readout learning and choice

During waking task epochs, *W*_rote_, *W*_knowledge_, and *b* were updated by a delta rule to predict reward. For scalar-value tasks, a state *i* with target reward *r*_*i*_ generated the prediction error

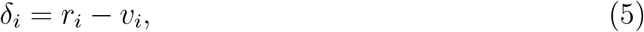

and the readout weights were updated as

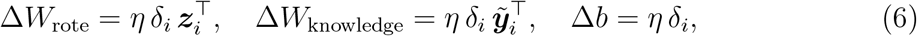

with *η* = 10^−4^.

For adjacent-pair transitive-inference training, the two candidate values in each pair were updated together using a pair-centered prediction error. For pair *p*, let ***r***_*p*_ and ***v***_*p*_ denote the two-dimensional target and predicted value vectors for the two candidate items in that pair. The pair-centered error was defined as

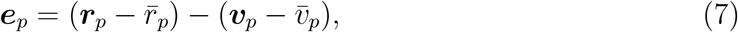

where 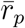 and 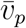 denote the mean target and predicted values within pair *p*, respectively. The same delta-rule update was then applied using ***e***_*p*_ in place of the scalar prediction error.

When no schema was available, the schema input to the task-value readout was set to zero, so that it carried no structured information. After a sleep period, a selected snapshot of *W*_schema_ was fixed, and task-value readout learning resumed using both the direct and schema-dependent pathways.

Choices were generated from value differences using a softmax policy with temperature *T* = 0.1. For binary tasks represented by a single scalar value *v*_*i*_, the action associated with *v*_*i*_ was compared against an alternative action with a reference logit of zero. The probability of choosing the action associated with *v*_*i*_ was

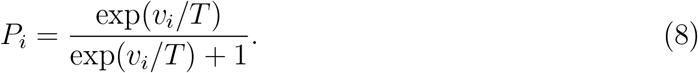

For transitive inference, each candidate item had its own value estimate, and the probability of choosing item *a* over item *b* was

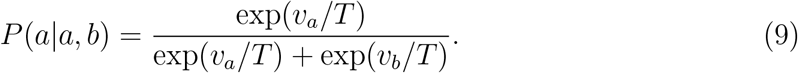

### Sleep-dependent schema learning

During sleep, *W*_schema_ was updated by Hebbian nonlinear dimensionality reduction [39], a biologically plausible learning rule that approximates the t-SNE [42] using a three-layer feedforward circuit and a three-factor Hebbian plasticity rule.

In this framework, repeated replay of experienced inputs in random order provides pairs of consecutive samples, allowing the circuit to infer input- and schema-space similarities from the activity patterns at times *t* − 1 and *t*. The sparse intermediate layer allows different states to be adjusted independently in the low-dimensional schema space. A global factor compares changes in input- and schema-space similarities and gates synaptic plasticity so that pairwise relationships among experienced states are preserved in the schema representation. The implementation follows Yoshida and Toyoizumi [39]; we summarize the variables used in the present study below.

At each sleep step, two successive replayed states (***x***_*t*−1_, ***x***_*t*_) were sampled in random order from the *N* state patterns, yielding the corresponding intermediate-layer activities (***z***_*t*−1_, ***z***_*t*_) and schema activities (***y***_*t*−1_, ***y***_*t*_). Here, ***y***_*t*_ denotes the raw schema activity 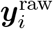 in the previous sections.

Input-space similarity between successive replays was estimated through gated axonal signals from the intermediate layer. For each intermediate unit *j*, the axonal activity *a*_*t,j*_ was set to one only when *z*_*t,j*_ was the largest positive response of that unit across all *N* state patterns, and was set to zero otherwise. This gating restricts *a*_*t,j*_ = 1 to the state pattern most strongly represented by unit *j*. The transmitter release at axon *j* was

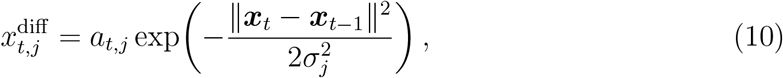

where *σ*_*j*_ is a synapse-specific scale parameter. The postsynaptic input-similarity signal was the sum of these contributions normalized by their time averages,

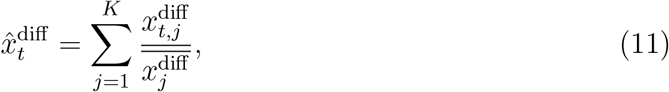

where *K* is the number of intermediate-layer units. This signal approximates the conditional similarity *p*_*i*|*j*_ used in t-SNE. This formulation corresponds to the input-specific normalization used in the original t-SNE.

Schema-space similarity was estimated with a Student-*t* kernel,

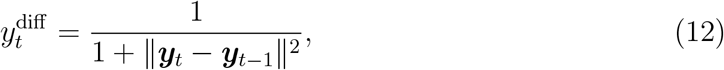

and its normalized counterpart,

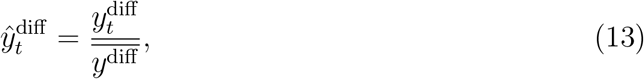

approximates the t-SNE output similarity *q*_*ij*_. The time averages 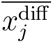 and 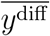 were maintained as running averages with time constant *τ* = 100.

The global factor was then defined as

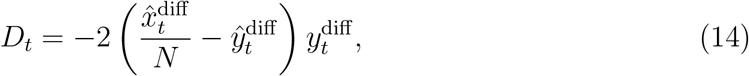

which compares input-space and schema-space similarities.

The synaptic update of *W*_schema_ was the product of this global factor, the change in postsynaptic schema activity, and the change in presynaptic intermediate-layer activity:

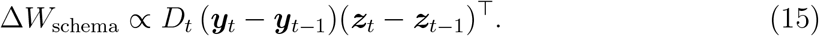

As shown in [39], the expected value of this three-factor Hebbian update approximates the negative gradient of the t-SNE objective, namely the KL divergence between input-space and output-space similarities, with respect to *W*_schema_.

The scale parameters *σ*_*j*_ were slowly adjusted so that the estimated perplexity approached a target value; see [39] for the update equations for *σ*_*j*_ and the associated entropy estimate. Following the Hebbian nonlinear dimensionality reduction implementation, the first 500 sleep epochs were used as a warm-up period for the similarity normalization and perplexity estimates, after which *W*_schema_ was updated. For numerical stability, synaptic updates were conducted using Adam [48] (*β*_1_ = 0.9, *β*_2_ = 0.999, learning rate 0.1), as in the original implementation [39].

### Manifold alignment for transfer

To model transfer across related tasks, we introduced a manifold-alignment term that attracted the current schema toward the schema of a previous task. Let 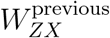and 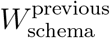 denote the fixed intermediate projection and schema weights from the previous task. For an input from the new task, ***x***_*i*_, the soft assignment to the previous task’s prototypes was

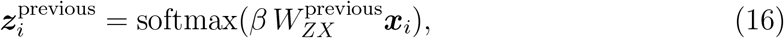

and the corresponding previous-schema coordinate was

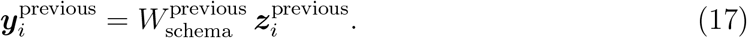

Alignment was applied only when this assignment to the previous task’s prototypes was sufficiently confident:

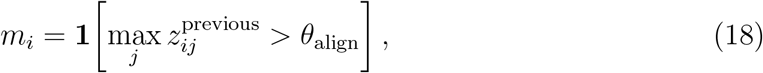

where 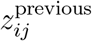 denotes the *j*-th component of 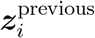 and *θ*_align_ = 0.1. For the current task, ***z***_*i*_ and ***y***_*i*_ denote the intermediate- and schema-layer activities for the input ***x***_*i*_. The alignment term added to the sleep update was

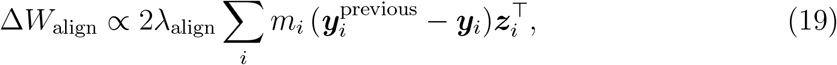

where *λ*_align_ = 10.0. The resulting sleep update therefore combined two constraints: preservation of the current task’s local relational structure and attraction toward the previous schema manifold.

### Spatial alternation task

The spatial task was a delayed alternation problem in which the agent had to alternate between left and right choices across successive traversals of a maze. The sensory input contained a place component and an action/reward-context component. For the hex maze, the place representation occupied a 10-dimensional subspace; for the square maze, it occupied a separate 6-dimensional subspace. Each vertex was assigned to one dimension, and points along an edge were represented by linear interpolation between the two endpoint vertices. The two mazes therefore used orthogonal place subspaces within a shared 16-dimensional place vector. An additional 6-dimensional action/reward-context vector encoded three binary variables near the decision point: the current left/right choice, the return direction from the previous trial, and the reward outcome at the choice point.

Place activity was temporally convolved with an exponential kernel (*r* = 0.99, window length 12) to provide short-term trajectory context. Small Gaussian noise with standard deviation 0.01 was added to the place subspace of the active maze. Each traversal contained 24 time steps in the hex maze and 16 time steps in the square maze. At each left/right decision, value was averaged over the four-step window ending at the branch point. The reward signal was non-zero only at decision-related time steps and indicated whether the current left or right choice was correct. Correct and incorrect choices were assigned rewards of 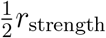 and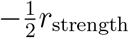, respectively, with *r*_strength_ = 2.5. Task-value readout updates were restricted to decision-related time steps; non-decision samples were excluded from the delta-rule update.

For learning in the hex maze, each independent simulation consisted of 160 epochs, with sleep inserted at epoch 150. Each wake epoch generated a trajectory containing eight choices, and the task-value readout was updated. Sleep consisted of 1000 epochs of Hebbian nonlinear dimensionality reduction with a target perplexity of 30. The dimensionality of the schema representation was set to three. For transfer to the square maze, the task-value readout weight *W*_knowledge_ and bias *b* learned in the hex maze were used to initialize square-maze learning. Only the aligned condition included the manifold-alignment term during sleep in the square maze. Schema untangling was quantified as the Euclidean distance between the mean schema coordinates of left- and right-choice decision states, computed over a four-step window ending at each branch point along a canonical trajectory. For evaluation of expected reward and schema untangling, we used a fixed noise-free canonical trajectory.

### Reversal learning task

We considered a probabilistic reversal learning task. The latent rule *h*_*t*_ ∈ {+1, −1} reversed every 10 trials in Task 1 and every 12 trials in Task 2. Task 1 used the stimulus pair *P/Q*, whereas Task 2 used a distinct pair, *R/S*. Under one rule, the first item in the current task (*P* in Task 1 and *R* in Task 2) was rewarded with probability 0.9; under the opposite rule, the second item (*Q* in Task 1 and *S* in Task 2) was rewarded with probability 0.9. The state contained the correct-choice history over the previous 9 turns, with each turn encoded as a two-dimensional one-hot vector. Task 1 and Task 2 histories were embedded in orthogonal subspaces, yielding a 36-dimensional task-history vector.

To provide a compact reward-history cue, the state was augmented with a 4-dimensional reward-related auxiliary vector. This vector was derived from an internal learner that tracked whether the first item in the current task was correct. At each trial *t*, a teaching signal *r*_*t*_ was set to +1 when the first item was correct and to −1 otherwise, and was used to update a running estimate 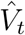 with learning rate 0.1. The resulting prediction error 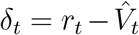 and its five-trial cumulative sum 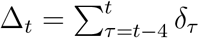 were used to define four one-hot indicator channels. The four channels were set to 1 only at the trials where *δ*_*t*_ reached its epoch-wise maximum or minimum and where Δ_*t*_ reached its epoch-wise maximum or minimum, and were 0 otherwise. The resulting input dimensionality was 40. Small Gaussian noise with standard deviation 10^−4^ was added to the history representation.

The task-value readout teaching signal was set to 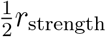 when the first item in the current task was correct and to 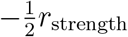 when it was incorrect, where *r*_strength_ = 2.5. Each simulation contained 15 wake epochs, with sleep inserted at epoch 5. Each wake epoch contained 100 turns in Task 1 and 120 turns in Task 2. The task-value readout was updated during each wake epoch. Sleep consisted of 1000 epochs of Hebbian nonlinear dimensionality reduction with a target perplexity of 30. The dimensionality of the schema representation was set to three. For transfer, the task-value readout weight *W*_knowledge_ and bias *b* learned in Task 1 were used to initialize Task 2. Only the aligned condition included the manifold-alignment term during sleep in Task 2.

### Transitive inference

The transitive inference task contained five items with latent order *A > B > C > D > E*. Training experience consisted only of adjacent pairs (AB, BC, CD, and DE). Each item was represented as an orthogonal vector. For each adjacent pair, the input specified the two presented items and the candidate choice, giving eight prototype choice states in a 10-dimensional input space. Each prototype was repeated 10 times with Gaussian noise (standard deviation 0.1), giving 80 training samples. The correct candidate in each adjacent pair had reward +1 and the incorrect candidate had reward −1.

Each independent simulation contained 10 epochs, with sleep inserted at epoch 5. Sleep consisted of 5000 epochs of Hebbian nonlinear dimensionality reduction with a target perplexity of 20. The dimensionality of the schema representation was set to two. Adjacent-pair performance was defined as

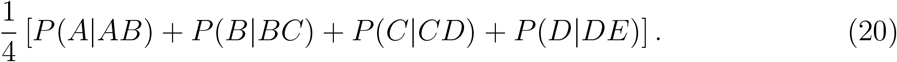

Relational generalization was assessed with non-adjacent probe pairs, including *P* (*B*|*BD*) and *P* (*A*|*AE*), without direct training on these pairs.

### Compositional transitive-inference and reversal-learning task

We constructed a composite network in which a transitive-inference network was followed by a reversal-learning network (Fig. 5b). The transitive-inference network had been pretrained on the *A*–*E* hierarchy as in Fig. 4, and the reversal-learning network had been pretrained independently in a two-item reversal task using stimuli *P/Q* that were distinct from the transitive-inference items. During target-task learning, the transitive-inference network was frozen and learning was restricted to the reversal-learning network (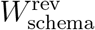,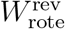,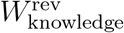, and the corresponding bias).

For each independent simulation, the transitive-inference network defined source values *V*_TI_(*ij, k*) for all pairs (*ij*) of the items {*A, B, C, D, E*} and the two candidate items *k* in each pair. Each *V*_TI_(*ij, k*) was the task-value readout output evaluated at the input state in which pair *ij* was presented and candidate *k* was queried. For the reversal-learning context input, these two source values were converted into a two-dimensional TI-derived choice-probability vector by applying the same softmax policy used for transitive-inference choice. The recent history input to the reversal-learning network consisted of these TI-derived choice-probability vectors, ordered so that the vector corresponded to the currently rewarded candidate under the latent reversal rule. Thus, the reversal-learning network did not receive item identities directly in the TI conditions; it received a history of TI-derived relational choice signals. In the without-transitive-inference condition, this TI-derived probability history was replaced by high-dimensional raw trial-state history encoding the presented pair and the rewarded candidate.

During adjacent-pair training, *V*_TI_ was used only for (*ij*) ∈ {*AB, BC, CD, DE*}. We compared three sources for these values: (i) *transitive inference after sleep*, in which task-value readout weights were trained using the final schema; (ii) *transitive inference before sleep*, in which task-value readout weights were trained until sleep; and (iii) *without transitive inference*, in which the transitive-inference source values were replaced by highdimensional inputs. When these inputs were provided to the reversal-learning network, recent correct choices were encoded as flattened vectors over the previous nine turns. Transitive-inference network training used schema dimensionality two, 5000 sleep epochs, and a target perplexity of 20.

The target task was a history-dependent reversal task over the four adjacent transitive-inference pairs. At each turn, one adjacent pair was sampled uniformly, and the latent ordering rule alternated every 10 turns between *A > B > C > D > E* and *A < B < C < D < E*. In the TI conditions, the task-value readout was trained on every turn, while in the *without transitive inference* condition the readout was updated only on AB-pair turns; all four pairs nevertheless contributed to the history input. Under the current rule, the higher-ranked item of the sampled pair was rewarded with probability 0.9. The history input described above (the TI-derived choice-probability vectors in the TI conditions, or the raw trial-state vectors in the without-transitive-inference condition) was embedded in task-specific subspaces and augmented with the same 4-dimensional reward-related auxiliary vector used in the reversal-learning task described above. Each simulation contained 15 epochs, with sleep inserted at epoch 5. Sleep consisted of 1000 epochs of Hebbian nonlinear dimensionality reduction with a target perplexity of 30, and the schema dimensionality was set to three.

The reversal-learning network produced a scalar context value *c*_*t*_ from its current schema and history state. For the presented pair *ij*, the transitive-inference-derived relational difference was

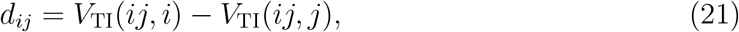

and the probability of choosing the first item was determined by the multiplicative composition of the reversal context and the transitive-inference-derived relational value:

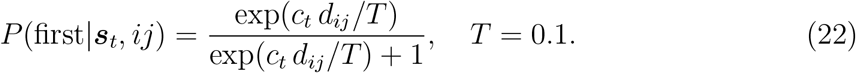

For each transitive-inference source, we further compared two alignment conditions in the reversal-learning network. In the *aligned* condition, sleep included the manifold-alignment term toward the previously learned *P/Q* reversal schema; in the *unaligned* condition, no alignment term was applied. Generalization to non-adjacent pair *BD* was evaluated by presenting all item pairs with the task-value readout weights frozen after adjacent-pair training. In the *only-alignment* condition (Fig. 5e), the reversal-learning network weights were taken immediately after the sleep period inserted at epoch 5, before any subsequent task-readout learning.

## Acknowledgments

We thank Tatsuya Haga, Takehiro Tottori, Kento Nakamura, Yoshihito Saito, and all members of the Toyoizumi and Inokuchi labs for helpful discussions. K.Y. was supported by JSPS KAKENHI Grant Numbers JP23K19415 and JP25K18569, and JST ACT-X Grant Number JPMJAX25LI. G.S. was supported by JSPS KAKENHI Grant Number JP23KJ0666. Y.K. was supported by JST ACT-X Grant Number JPMJAX25CA and JST BOOST Grant Number JPMJBS2418. K.I. was supported by JSPS KAKENHI Grant Number JP23H05476 and JST CREST Grant Number JPMJCR23N2. T.T. was supported by RIKEN Center for Brain Science, RIKEN TRIP initiative (RIKEN Quantum), JST CREST Grant Number JPMJCR23N2, and JSPS KAKENHI Grant Number JP25K24466.

## Author contributions

K.Y. and T.T. conceived the project and developed the main theoretical framework through discussion. K.Y. conducted the numerical simulations. G.S. and Y.K. contributed to discussions on model refinement. K.I. contributed to project development from the perspective of experimental neuroscience. K.Y. and T.T. wrote the manuscript. All authors reviewed and edited the manuscript.

## Data availability

No new experimental data were generated in this study.

## Code availability

The source code will be made available in a public repository upon publication.

## Competing interests

The authors declare no competing interests.

## Notes

### Competing Interest Statement

The authors have declared no competing interest.

